# Phosphatidylinositol 4-kinase III beta regulates cell shape, migration and Focal Adhesion number

**DOI:** 10.1101/828046

**Authors:** Patricia Bilodeau, Daniel Jacobsen, Denise Law-Vinh, Jonathan M Lee

**Author notes:** These authors contributed equally to this work and are listed alphabetically (Patricia Bilodeau, Daniel Jacobsen, Denise Law-Vinh). **Corresponding Author:** Jonathan M Lee. Department of Biochemistry, Microbiology, and Immunology, University of Ottawa, 451 Smyth Road, Ottawa, ON K1H 8M5, Canada.

## Abstract

Cell shape is regulated by cell adhesion and cytoskeletal and membrane dynamics. Cell shape, adhesion and motility have a complex relationship and understanding them is important in understanding developmental patterning and embryogenesis. Here we show that the lipid kinase phosphatidylinositol 4-kinase III beta (PI4KIIIβ) regulates cell shape, migration and Focal Adhesion (FA) number. PI4KIIIβ generates phosphatidylinositol 4-phosphate from phosphatidylinositol and is highly expressed in a subset of human breast cancers. PI4KIIIβ and the PI4P it generates regulate a variety of cellular functions, ranging from control of Golgi structure, fly fertility and Akt signaling. Here, we show that loss of PI4KIIIβ expression decreases cell migration and alters cell shape in NIH3T3 fibroblasts. The changes are accompanied by an increase in the number of FA in cells lacking PI4KIIIβ. Furthermore, we find that PI4P-containing vesicles move to the migratory leading-edge during migration and that some of these vesicles tether to and fuse with FA. Fusion is associated with FA disassembly. This suggests a novel regulatory role for PI4KIIIβ and PI4P in cell adhesion and cell shape maintenance.

## Introduction

A culture of genetically identical NIH3T3 fibroblasts displays a striking visual diversity. A single microscopic field of view is populated with elongated cells, round cells and yet others with extraordinarily complex geometries. Furthermore, some NIH3T3 cells are solitary while others cluster themselves into multicellular groups. Lastly, some fibroblasts are stationary while others are motile. These moving cells display marked differences in speed and directionality. This fascinating architectural and behavioral diversity is at the root of processes such as organismal development and patterning.

Cell shape has important implications in cell function (Bellas and Chen, 2014; Gilbert and Weaver, 2017). For example, the spreading of a cell in two-dimensional culture regulates both sensitivity to apoptosis and proliferative capacity (Chen *et al*., 1997). The spreading and shape of a cell results from a complex and dynamic interaction between its cytoskeleton, the plasma membrane and the extracellular matrix (ECM) (Mogilner and Keren, 2009). In fish keratinocytes, which assume a hemi-spherical appearance in 2-D culture, cell shape is determined by a balance between the protrusive force of actin polymerization and tension in the plasma membrane (Keren *et al*., 2008). In other cells, cell shape is regulated by lipid constituents of the plasma membrane (Wen *et al*., 2018) and by membrane-binding proteins that induce curvature (Nishimura *et al*., 2018). In migratory fibroblasts, which assume a wedge-shaped form, cell shape is additionally regulated by the presence and strength of adhesive structures at the cell periphery (Satulovsky *et al*., 2008). These adhesive structures provide sites for attachment to the ECM and the sensing of its mechanical properties. Cellular adhesions also serve as sites for the generation of contractile forces (Bosch-Fortea and Martin-Belmonte, 2018).

Perhaps the most important of mammalian adhesive structures are Focal Adhesions (FA) (Burridge, 2017). FA are oval-shaped, multi-protein complexes, generally 2 μm wide and 3-10 μm long, couplind the actin cytoskeleton to the ECM via transmembrane integrin heterodimeric receptors (Parsons *et al*., 2010). During cell migration, FA are created from smaller integrin-containing structures described as either Focal Complexes or Nascent Adhesions (Parsons *et al*., 2010). In a mature FA, the cytoplasmic tails of the integrin molecules are linked to actin filaments via adapter proteins such as talin, paxillin, zyxin and vinculin (Kanchanawong *et al*., 2010). FA are sites of mechano-transduction and activate a diverse array of signal transduction pathways (Geiger *et al*., 2009). FA number and size are critical in the maintenance of migratory capacity (Kim and Wirtz, 2013a), cell spreading (Kim and Wirtz, 2013b) and cell shape (Chen *et al*., 2003; Mogilner and Keren, 2009).

In this report, we find that the lipid kinase PI4KIIIβ has an important role in regulating cell shape, migration and FA number. A Golgi-resident enzyme, PI4KIIIβ is one of four mammalian proteins that generate PI4P from PI (Balla, 1998, 2013). PI4 kinases have been implicated in several human cancers (Waugh, 2012). PI4KIIIβ is a likely human oncogene based on its high expression in a subset of human breast tumours and its ability to co-operate with the Rab11a GTPase to activate signaling through Akt (Morrow *et al*., 2014). PI4KIIIβ and its homologues have multiple physiological roles, ranging from controlling Golgi structure in yeast to regulating male fertility and development in Drosophila (Godi *et al*., 1999; Polevoy *et al*., 2009). PI4KIIIβ also regulates *in vitro* morphogenesis of human breast cells (Pinke and Lee, 2011).

## Results

### PI4P vesicles move to the migratory leading edge

To explore a role for PI4P in cell migration, we used two PI4P biosensors, GFP-FAPP1 and GFP-P4M, to study intracellular PI4P localization during NIH3T3 migration. The GFP-FAPP1 biosensor consists of GFP conjugated to the PH (Pleckstrin Homology) domain of FAPP1 (four-phosphate-adaptor protein 1)(Balla, 2007). The GFP-P4M biosensor consists of GFP conjugated to a single P4M domain containing residues 546–647 of the *L. pneumophila* SidM protein(Hammond *et al*., 2014). Both biosensors bind to PI4P directly. However, FAPP1 binds to PI4P only when complexed to the Arf-1 GTPase (Balla *et al*., 2005).

In migrating NIH3T3 cells, FAPP1 shows a broadly perinuclear distribution consistent with Golgi localization (Fig 1A, Supplementary video S1). Large (0.8 – 1.6 um diameter) and small vesicles (0.2 – 0.45 um diameter) are visible. The large vesicles (lv) are predominantly perinuclear while the small vesicles (sv) localize to the cell periphery. P4M shows a somewhat similar localization pattern with many large vesicles (lv) in the perinuclear region. Different from FAPP1, many of these perinuclear vesicles appear hollow (hv) (Fig 1A, Supplementary video S2). As is the case with FAPP1, there are small vesicles in the cell periphery but these seem to be much more numerous in P4M transfected cells. In direct contrast to FAPP1 and as observed previously in COS7 cells (Hammond *et al*., 2014), the P4M reporter localizes to plasma membrane structures. Some plasma membrane features at the migratory leading edge features are reminiscent of ruffles (r), actin rich structures that move away from the leading edge and derive from poor lamellar adhesion to the growth substrate (Borm *et al*., 2005). The difference between FAPP1 and P4M staining presumably reflects the requirement that FAPP1 interact with Arf-1 associated PI4P pools.

**Figure 1.**
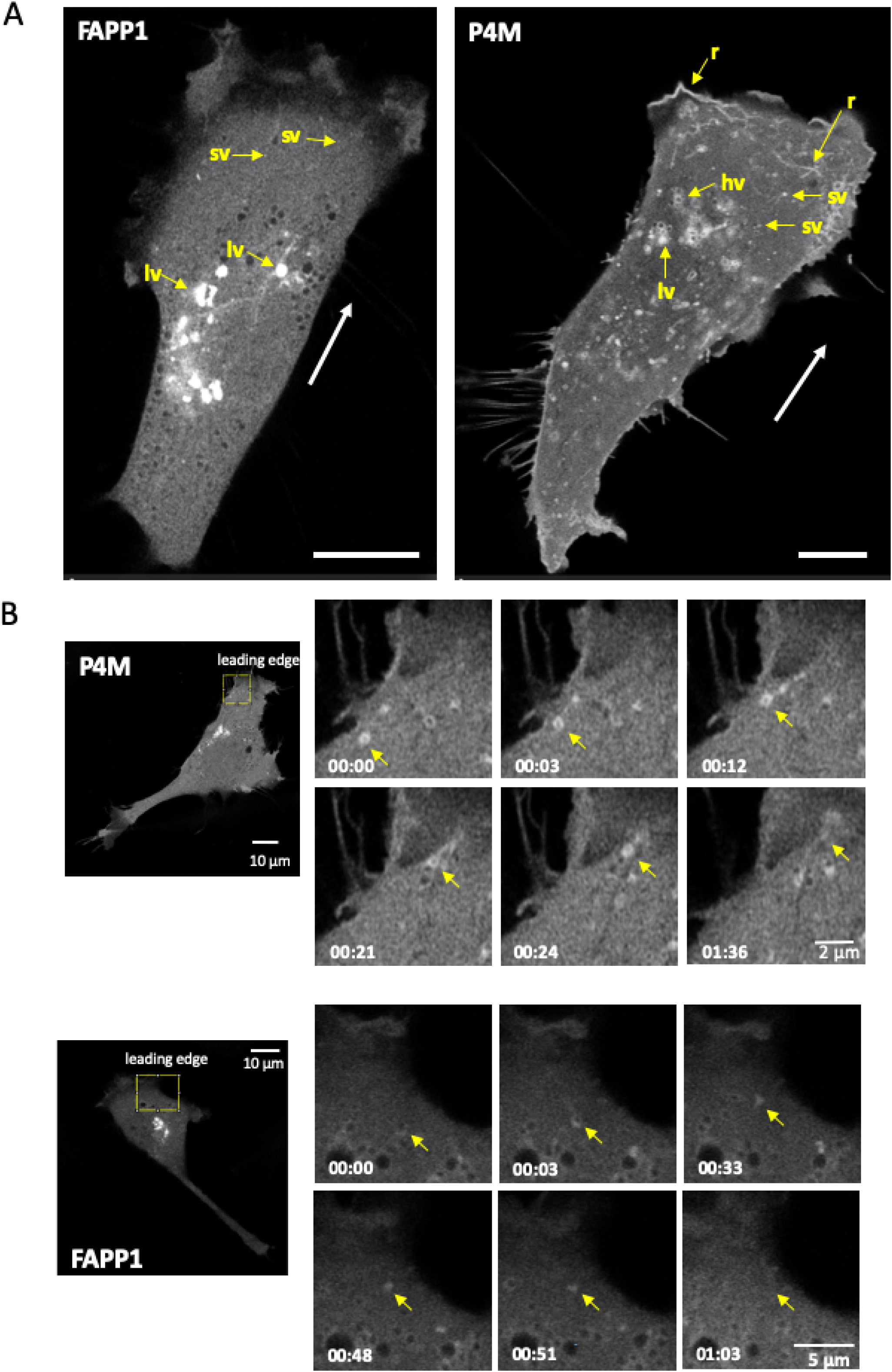
PI4P containing vesicles move to and fuse with the migratory leading edge. **A)** PI4P-contining vesicles are visualized in NIH3T3 fibroblasts undergoing migration (direction indicated by white arrow) using FAPP1 and P4M reporters. In FAPP1 cells, large and small vesicles, labelled lv and sv respectively, are seen. In P4M cells, hollow vesicles (hv) and linear structures reminiscent of ruffles can also be seen. Scale bar is 10 μm. **B)** PI4P-contining vesicles in motile cells move to and fuse with the leading edge. An individual vesicle is identified by the yellow arrow. Elapsed time and scale bars are indicated.

During migration, the large vesicles of both P4M and FAPP1 show only limited intracellular movement and remain generally perinuclear (Supplementary videos S1 & S2). In contrast, the peripheral small vesicles show a high degree of mobility (Supplementary videos S1 & S2). In addition, we observe multiple P4M and FAPPI small vesicles moving to the migratory leading edge and then disappearing (Figure 1B, Supplementary videos S3 & S4). This is consistent with vesicular delivery of PI4P-containing vesicles, both Arf-1 associated and Arf-1 free forms, to the plasma membrane. The absence of FAPP1 staining at the plasma membrane suggests that once the vesicle delivers the PI4P/Arf-1 cargo, either Arf-1 dissociates or PI4P is converted by PI4P kinases to another PI. In the case of P4M, the persistence of PI4P in some parts of the leading edge indicate that some of the membrane delivered PI4P remains as PI4P. In addition to their movement to the plasma membrane, many small P4M and FAPP1 vesicles show considerable directional freedom. Both retrograde and anterograde movements are observed (Supplementary Videos S3 & S4). However, the movement of small vesicles is generally toward the migratory leading edge.

The movement of vesicles towards the migratory leading edge is mirrored in the visibly unequal intracellular distribution of both FAPP1 and P4M vesicles in the migratory leading edge compared to the trailing edge. To further explore this, we used automated image analysis to quantify the distribution of PI4P vesicles in migratory and non-migratory cells (Supplementary Videos S5, S6, S7 & S8 show representative cells). In migratory cells, we counted vesicles in both the leading and trailing edges. In stationary cells, we counted vesicles present in two opposing edges of the cell. In stationary cells there are a generally an equal number of P4M and FAPP1 marked PI4P vesicles at two opposite edges (Fig 2B, open circles; Supplementary Videos S6 & S8). On the other hand, migratory cells show a greater number of vesicles at their leading edge compared to the trailing edge (Fig 2B, closed; Supplementary Videos S5 & S7). This is consistent with the idea that the transport of PI4P to the migratory leading edge is an part of the directional cell migration machinery.

**Figure 2.**
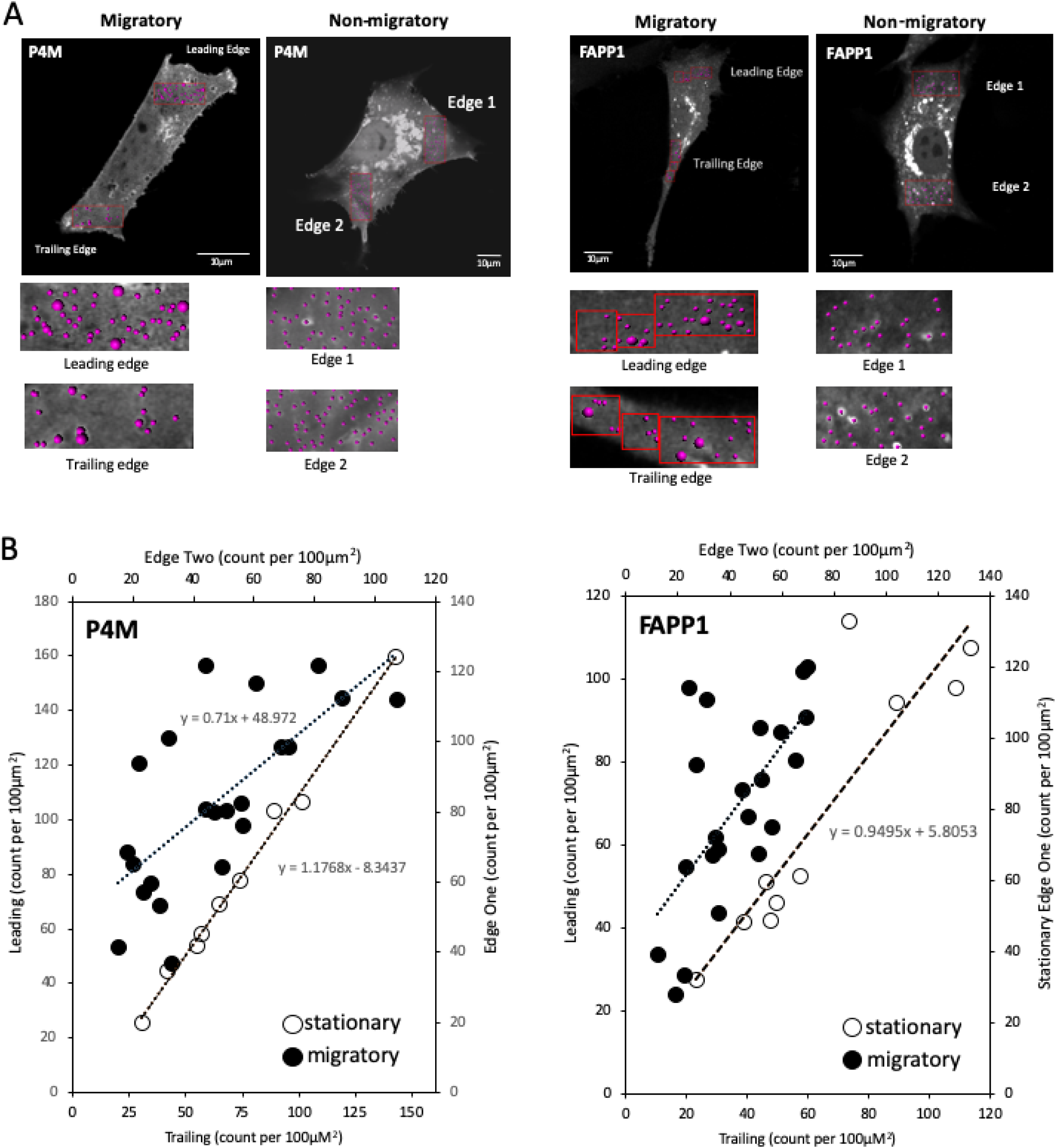
PI4P containing vesicles have polarized distribution in migrating cells. **A)** PI4P-contining vesicles are visualized in NIH3T3 fibroblasts that are either migratory (leading and trailing edges indicated) or non-migratory. Vesicles are counted by volume rendering in. **B)** The number of PI4P vesicles, either P4M or FAPP1 labelled, at the leading edge is plotted as a function of the vesicles at the trailing edge in migratory cells (•). For non-migratory cells, the number of vesicles is plotted for two opposite edges (o). The line of best fit is shown.

### PI4KIIIβ deletion decreases cell migration

To explore a role for PI4P in cell migration, we used CRISPR to delete the PI4KIIIβ gene in mouse NIH3T3 fibroblasts. PI4KIIIβ generates PI4P in the Golgi (Balla and Balla, 2006; Balla, 2013). We then tested two independent PI4KIIIβ null lines for migratory capacity in wound healing assays. As shown in Fig 3A (left panel), the two CRISPR lines close a wound much slower that parental cells. Wild type NIH3T3 cells close half the wound in ∼18 hours, while the null lines take ∼25 hours. This defect in wound closure is rescued by re-expression of wild-type PI4KIIIβ (Fig 3A, middle panel).

**Figure 3.**
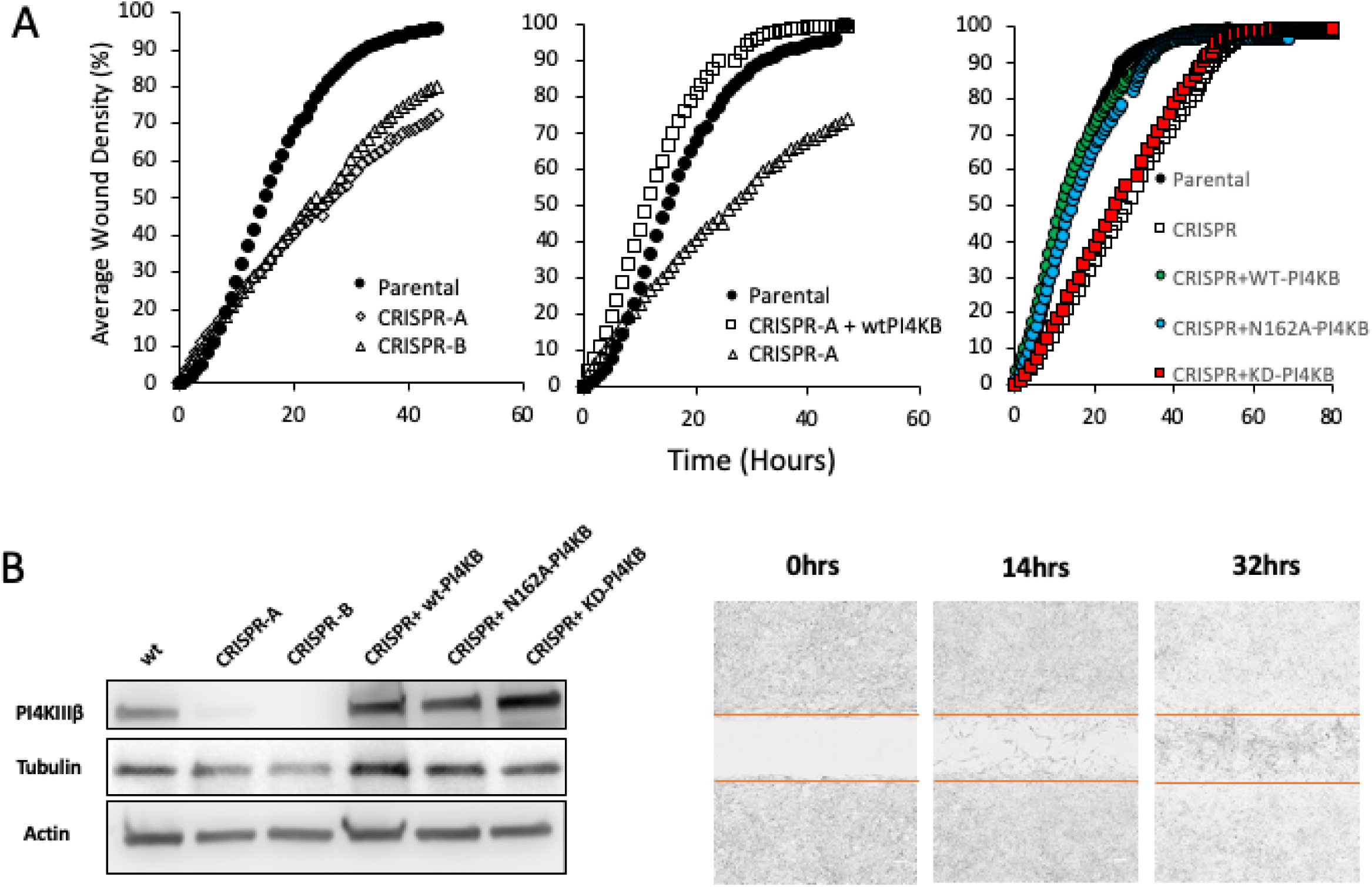
Loss of PI4KIIIβ impairs migration. **A)** (*Left panel*) The ability to close an *in vitro* wound is shown in two independent lines of PI4KIIIβ–deleted compared to parental cells. (*Centre panel*) Attenuated wound healing is rescued in the CRISPR line by re-expression of wild type PI4KIIIβ (wtPI4KB). (*Right panel*) Attenuated wound healing is rescued in the CRISPR line by re-expression of wild type PI4KIIIβ (WT-PI4KB) and kinase dead (KD-PI4KB) but not a Rab11-binding mutant (N162A-PI4KB). Experiments are representative of triplicates. **B)** (*Left panel*) Protein expression of PI4KIIIβ and tubulin and actin control in the cell lines. (*Right panel*) Inset shows a representative wound closure experiment. Scale bar in white is 30 μm.

The best characterized function of PI4KIIIβ is the generation of PI4P from PI (Balla, 1998). However, PI4KIIIβ also interacts with the Rab11a GTPase and is part of Rab11a-dependent pathway controlling endosome function (de Graaf *et al*., 2004; Polevoy *et al*., 2009; Burke *et al*., 2014). To determine which of these functions were involved in migration regulation, we expressed wt-PI4KIIIβ, kinase dead PI4KIIIβ (KD-PI4KIIB) (Zhao *et al*., 2000) or a PI4KIIIβ mutant that does not interact with Rab11a (N162A)(Burke *et al*., 2014) in a CRISPR line. These lines were then tested for migratory capacity. As shown in Figure 3A (right panel), a CRIPSR line expressing wt-PI4KIIIβ or the Rab11a-binding mutant N162A had similar closure times to those of parental fibroblasts. On the other hand, KD-PI4KIIIβ was unable to rescue would closure kinetics. This indicates that the ability to generate PI4P, rather than Rab11a interaction, is required PI4KIIIβ-mediated control of migration. Figure 3B shows protein expression of the PI4KIIIβ rescued cell lines.

### PI4KIIIβ regulates stress fiber appearance

In yeast, the PI4KIIIβ homolog Pik1 is required for maintaining Golgi structure (Walch-Solimena and Novick, 1999; de Graaf *et al*., 2004). Because cell migration involves repositioning of the Golgi between the nucleus and leading edge (Kupfer *et al*., 1982) and functional alterations of the Golgi affect migration (Xing *et al*., 2016; Ahat *et al*., 2019), we next investigated whether or not PI4KIIIβ depletion affected Golgi appearance. When stained for markers of either the cis-or trans-Golgi (GM130 and TGN46 respectively), NIH3T3 cells lacking PI4KIIIβ have Golgi that are visibly indistinguishable from wild type cells (Fig 4A). Moreover, in scratch wound assays PI4KIIIβ depletion does not visibly affect Golgi positioning to face the wound (Fig 4B). Microtubule structure also appears normal as does centrosome alignment away from the wound (Fig 4B). One the other hand, PI4KIIIβ depletion causes an increase in the number of cells with numerous stress fibers (Fig 4C). In a population of normal NIH3T3 cells, most cells show few stress fibers with only 3.1 ±5% of the population having more than 6 fibers per cells. On the other hand, the PI4KIIIβ depleted lines have a much larger fraction (11.2 ±8% and 18.9 ±13%) in their population with high stress fiber content

**Figure 4.**
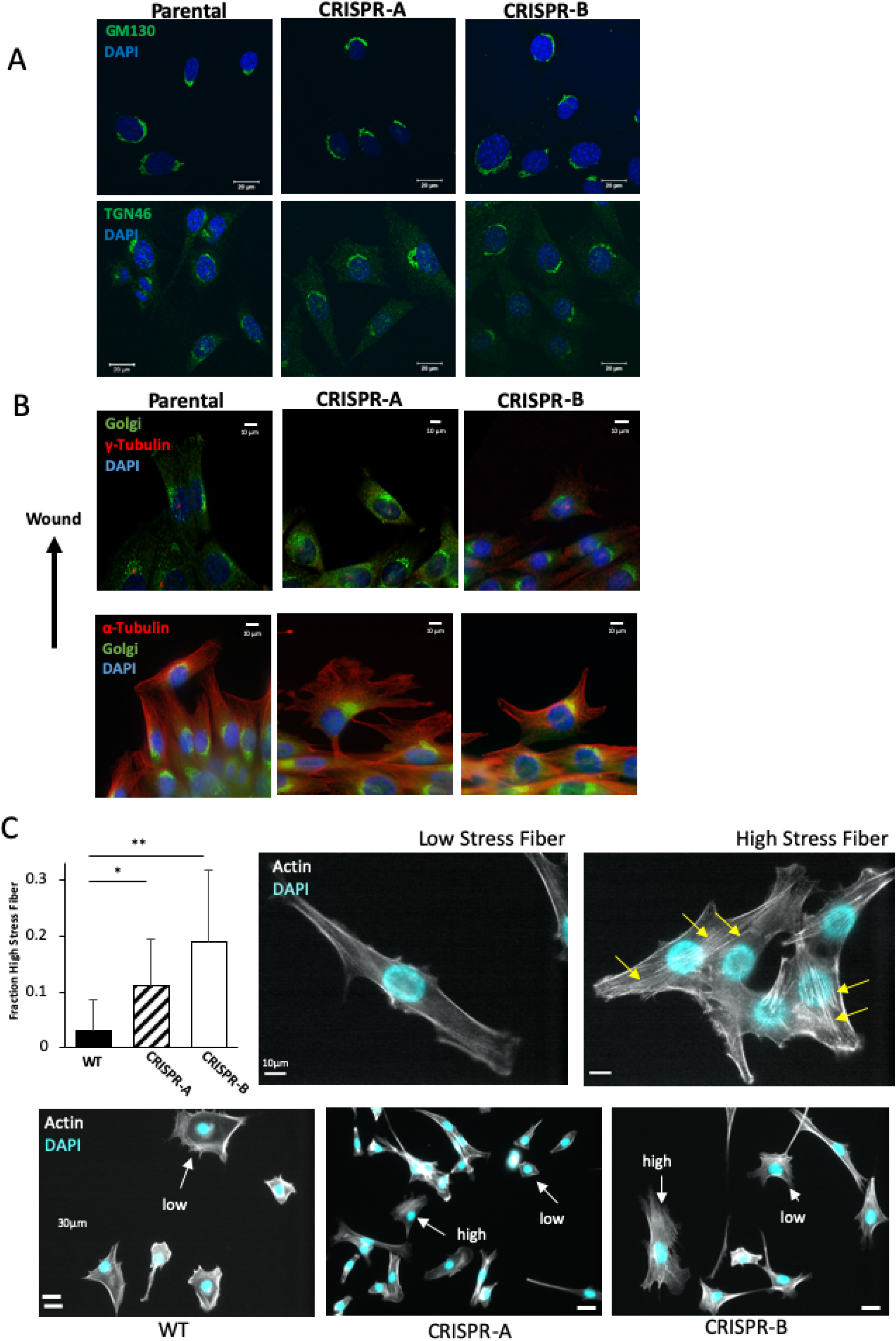
Loss of PI4KIIIβ alters stress fiber appearance. **A)** Cis- and trans-Golgi structure as visualized by GM130 and TGN46 staining respectively in wild type NIH3T3 cells and two independent lines of PI4KIIIβ-deleted cells. **B)** Golgi, centrosome (γ-tubulin) and microtubule (α-tubulin) structure visualized in cells entering a scratch wound. **C)** CRISPR cell lines both show significantly (t-test, p< 0.0001) more cells with high numbers of stress fibers compared to wild type cells. Representative cells with respectively low or high numbers of stress fibers (yellow arrows) are shown in the right panels. Bottom panels show representative fields of wild type and CRISPR lines showing cells with high or low stress fiber content.

### PI4KIIIβ regulates cell shape

In our studies of the cell cytoskeleton and cell migration, we noticed that cultures of the CRIPSR deleted NIH3T3 cells had a very different morphological appearance under phase than either parental or rescued cells. In our experience, most cultured NIH3T3 cells assume one of three broad shapes. The first is an elongated form (Fig 5A) and approximately half of wild type NIH3T3 cells assume this shape (Fig 5A Right Panel). The second most common shape is what we term “multidirectional”. These cells are roughly rectangular in shape with multiple pseudopodial protrusions. Approximately 25% of wild type cells are of this type (Fig 5A Right Panel). The remaining cells, which have a smaller, generally spherical appearance, we classified as “other”. The loss of PI4KIIIβ leads to a redistribution of cell shapes in both of the CRISPR lines. In freely migrating conditions, the number of elongated cells in the PI4KIIIβ deleted cells decreases by almost 50% and the number of multidirectional cells more than doubles (Fig 5A Top Panel). Similarly, cells present in the wound of a scratch migration assay show similar increase in the number of multidirectional cells and a decrease in elongated ones (Fig 5A Bottom Panel). Representative fields are shown in Fig 5B. As is the case with the wound healing assay, wt-PI4KIIB and the Rab11-binding mutant (N162A) were able to restore wild type shape distribution to the CRISPR lines, while the kinase dead PI4KIIIβ did not (Fig 5C). Parental cells and CRISPR lines rescued with either wild type PI4KIIB or N162A had 45-50% of cells as elongated, while the kinase dead rescued cells had nearly 35% as multidirectional, similar to the original CRSPR line. Similarly, parental cells and CRISPR lines rescued with either wild type PI4KIIIβ or N162A had 25-35% of cells as multidirectional, while the CRISPR and kinase dead rescued cells had nearly 50% as multidirectional. Thus, cell shape control by PI4KIIIβ, like wound healing, is dependent on PI4P generation rather than Rab11a interaction.

**Figure 5.**
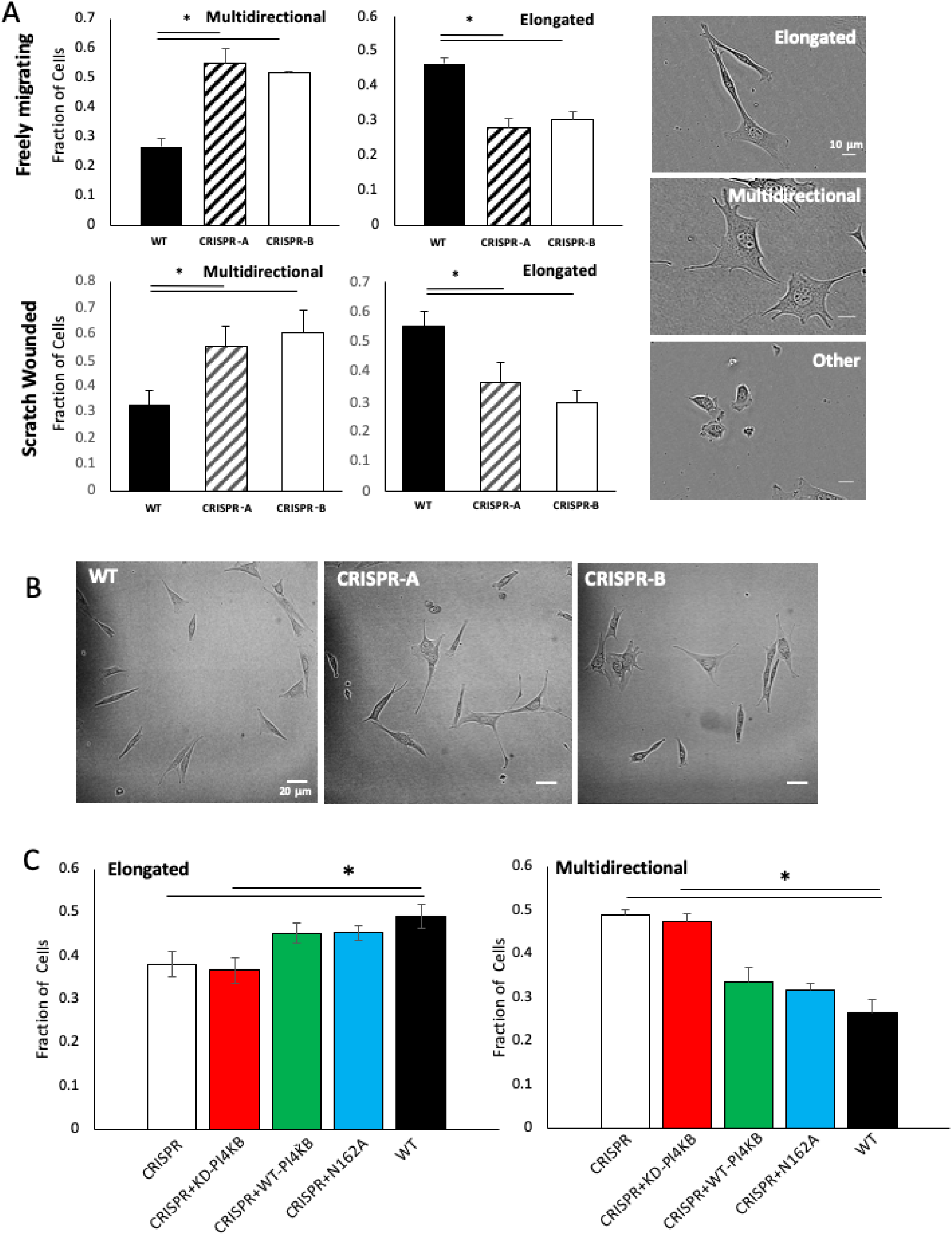
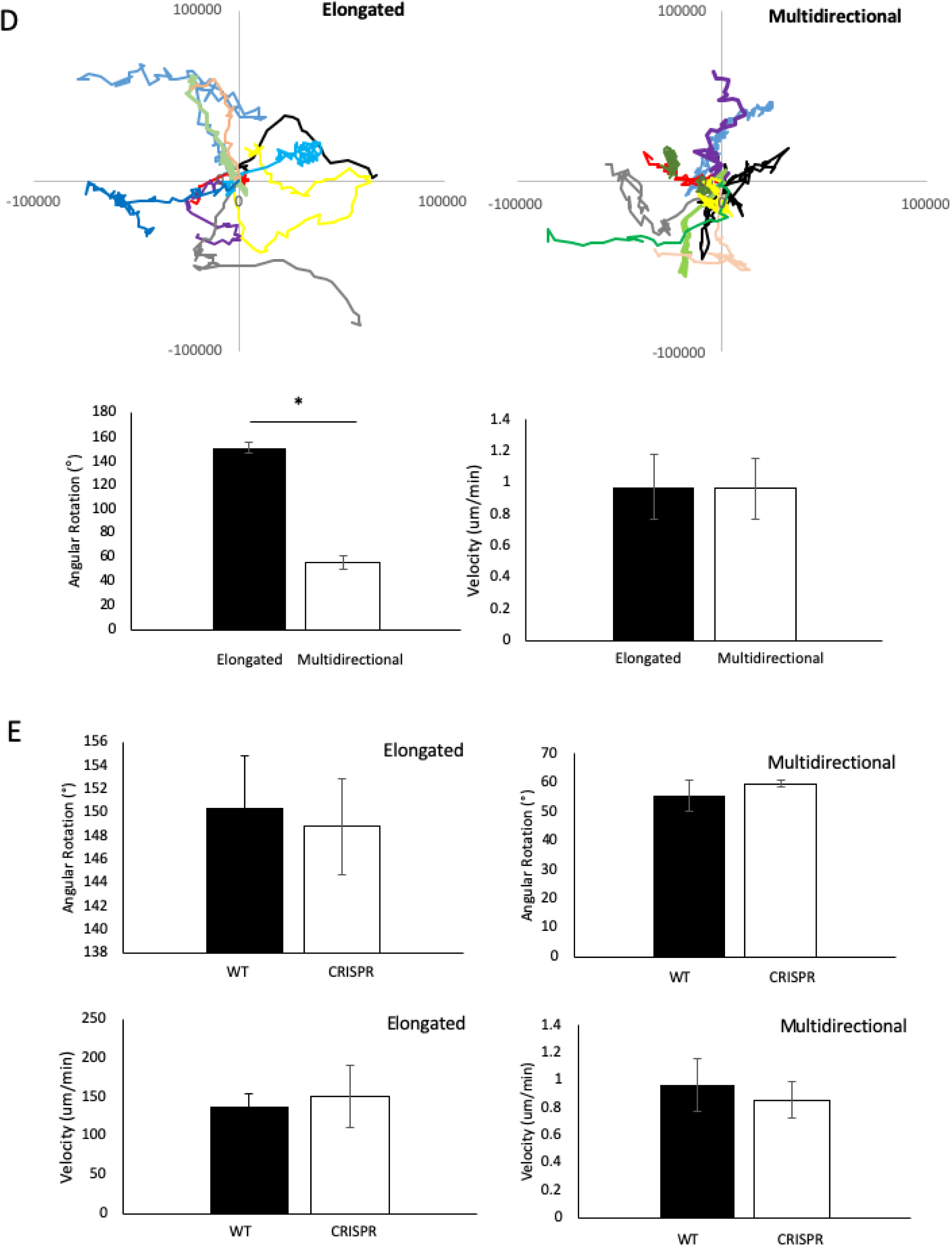
Loss of PI4KIIIβ alters cell shape distribution. **A)** Two independent lines of PI4KIIIβ - deleted cells have different population shape distributions (depicted in right panels) in both freely migrating conditions and in the wound of a scratch. The fraction of elongated cells in either line of CRISPR cells is significantly (p<0.001, t-test) lower than in a population of wild type cells. Similarly, the fraction of elongated cells in either line of CRISPR cells is significantly (p<0.001, t-test) higher than in a population of wild type cells. Results are the mean and standard deviation of triplicate independent measurements of at least 200 cells each from a minimum of 20 randomly selected fields **B)** Representative fields of each cell type. Scale bar is 20 μm. **C)** Re-expression of either wild-type- or N162A-PI4KIIIβ restores elongated and multidirectional shape distribution to the CRISPR cells. Expression of kinase dead PI4KIIIβ does not rescue cell shape changes and their elongated of multidirectional composition is similar to the CRSPR control and significantly different from wild type cells (p<0.05, t-test). Results are the mean and standard deviation of quadruplicate independent measurements of at least 200 cells from at least 20 randomly selected fields **D)** Distance travelled over time was monitored for 50 wild type NIH3T3 cells with either elongated or multidirectional morphologies. Tracks for 10 cells of each type are shown. Multidirectional cells have a significantly reduced average angular rotation that elongated cells (p<0.05, t-test). **E)** Quantification of migration patterns in wild type cells with either multidirectional or elongated morphologies. Mean angular rotation or velocity is presented as the mean and standard deviation of 50 cells collected over at least 30 minutes. Angular rotation is significantly lower in the multidirectional population compared to the elongated one (p< 0.05,t-test). Comparison between Angular Rotation and Velocity between wild type and CRISPR lines. Mean angular rotation or velocity is presented as the mean and standard deviation of 50 cells collected over at least 30 minutes.

We hypothesized that this change in population cell shape was related to migratory defects in the CRISPR knockout. As shown in Fig 5D, elongated wild type NIH3T3 cells exhibit different migratory behaviour that the multidirectional ones. Multidirectional cells tend to make sharp and frequent turns, while elongated cells oscillate back and forth only a more or less straight line (Fig 5D). This means that although their migratory velocity is similar (Fig 5D), elongated cells travel a further distance from their origin than their multidirectional counterparts (Fig 5D). The behaviour of individual CRIPSR cells, in either elongated or multidirectional shape classes, is similar to their wild type counterparts in that they have similar turning behaviours and recorded velocities (Fig 5E). This suggests that the loss of PI4KIIIβ is not affecting the migratory machinery per se. Rather it is affecting cell shape, which changes the migratory behavior of the cell population.

### PI4KIIIβ regulates Focal Adhesions

In order to further explore the regulation of cell shape and migration by PI4KIIIβ, we investigated what cellular structures might be interacting with PI4P-containing vesicles. We reasoned that Focal Adhesions (FA) might be one such structure since FA-dependent adhesion is likely to be involved in both cell shape, migration control and stress fiber formation (Chen *et al*., 2003; Kim and Wirtz, 2013a). To investigate possible FA and PI4P interaction during cell migration, we simultaneously imaged them in migrating cells using fluorescent Talin and P4M reporters. At the migratory leading edge (Fig 6A, Supplementary Video S9), PI4P vesicles can be observed moving to FA. Vesicles dock with FA and then disappear, presumably delivering their cargo to FA and the PI4P becoming metabolized. Similar behavior occurs at FA of the trailing edge (Fig 6B, Supplementary Video S10).

**Figure 6.**
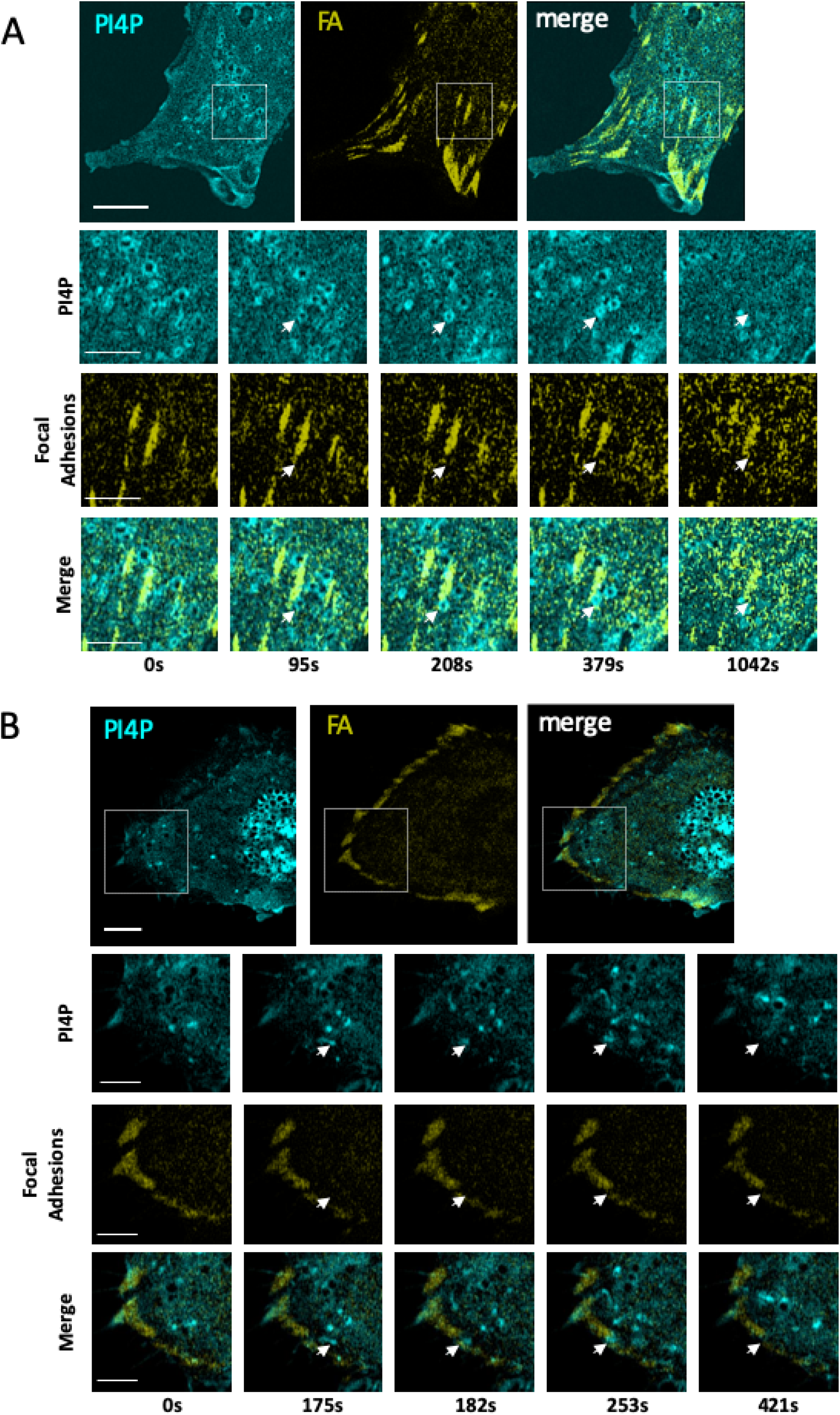
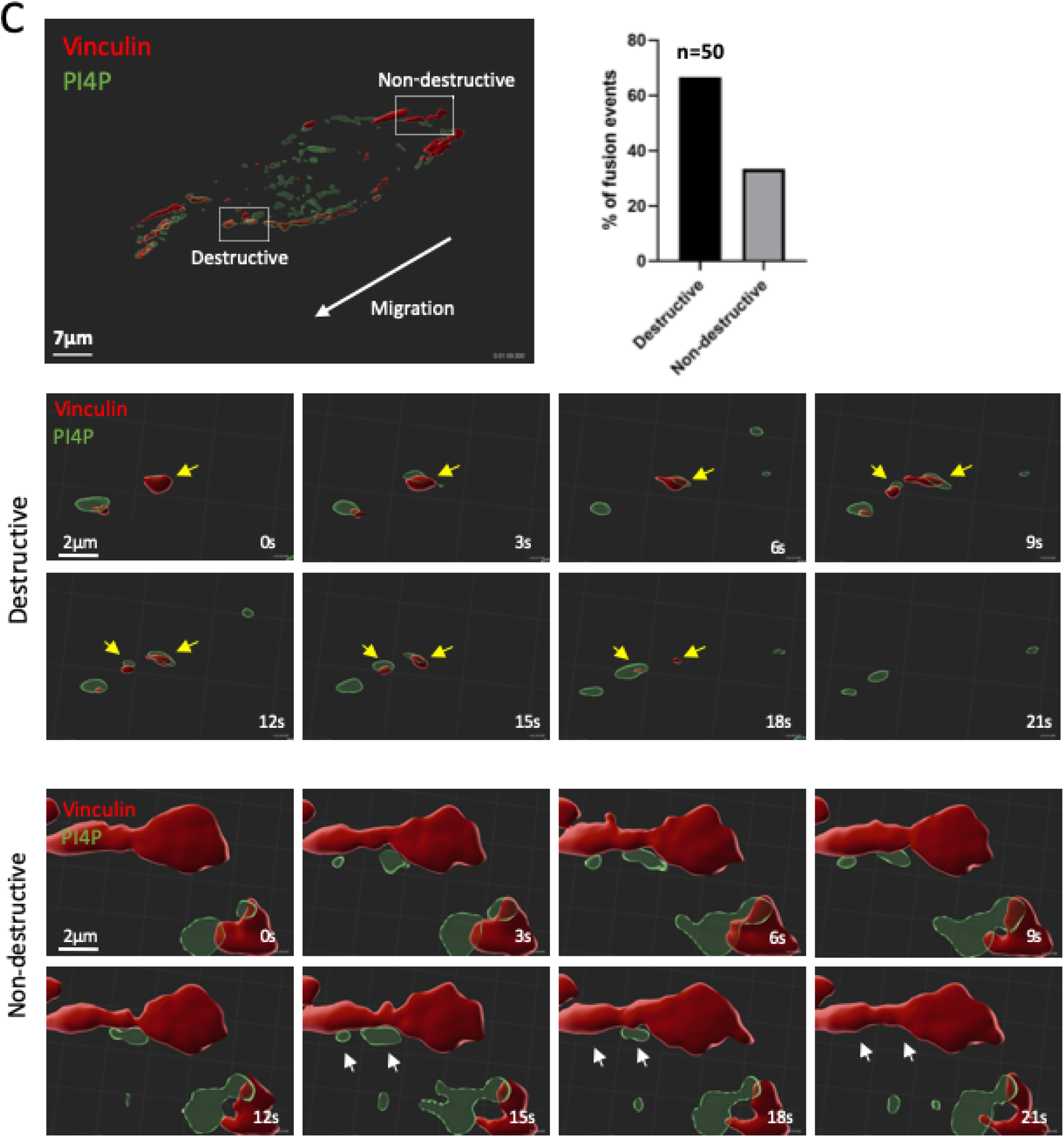
PI4P Vesicles move to and fuse with Focal Adhesions. **A)** Visualization of PI4P and Focal Adhesions (FA) in the leading edge of a migrating NIH3T3 cells. The time series indicates a single vesicle observed from the boxed inset. The fusing vesicle (red arrow) appears at 95s. Scale bar in the large image is 10 μm and is 3 μm in the enlarged images. **B)** Visualization of PI4P and FA in the trailing edge of a migrating NIH3T3 cells. The time series indicates a single vesicle seen from the boxed inset. The fusing vesicle (red arrow) appears at 175s. Scale bar in the large image is 10 μm and is 3 μm in the enlarged images. **C)** Bessel Lattice Light Sheet imaging of PI4P vesicles (green P4M reporter) fusing with FA (red Vinculin reporter) in a migrating cell. Histogram shows that the majority of fusions events (n=50) cause FA destruction within 3-6s. Lower panels show enlarged fields of the upper panel where two fusion events, one causing FA destruction and the other not, can be seen.

To further explore delivery of PI4P vesicles to FA, we used a Bessel Lattice Light Sheet microscope (BLLSM) to image migrating NIH3T3 cells. The BLLSM illuminates a sample using a thin structured illumination beam and allows for long-term imaging with high axial and temporal resolution and with minimal photobleaching (Planchon *et al*., 2011). Using the BLLSM, we visualized 50 PI4P vesicle-FA fusion events in five independent migrating wild type NIH3T3 cells at 3s/volume. Cells were observed for ∼25 minutes each. As shown in Fig 6C, approximate two thirds of fusion events were associated with disappearance of the FA within 3-6 seconds. This suggests that PI4P or cargo in PI4P vesicles is involved in FA disassembly. To test this idea, we used an automated image analysis program to count FA in wild type and CRISPR deleted NIH3T3 cells (Horzum *et al*., 2014). As shown in Fig 7A, wild type NIH3T3 cells (n=60) had an average of 56.7±17 FA per cell. CRISPR deleted cells (n=59), on the other hand, had significantly (t-test, p<1 × 10^−5^) more FA, averaging 84.6±42 per cell. Rescue of the CRISPR line by PI4KIIIβ re-expression returned FA number to wild type levels (58.4±24). Thus, the number of FA/cell is dependent, at least in part, on PI4KIIIβ. In addition, FA in the CRISPR lines show defects in FAK phosphorylation in response to hypotonic stress (400 mM sucrose). Thus, loss of PI4KIIIβ allows for more FA per cell but these have reduced functionality.

**Figure 7.**
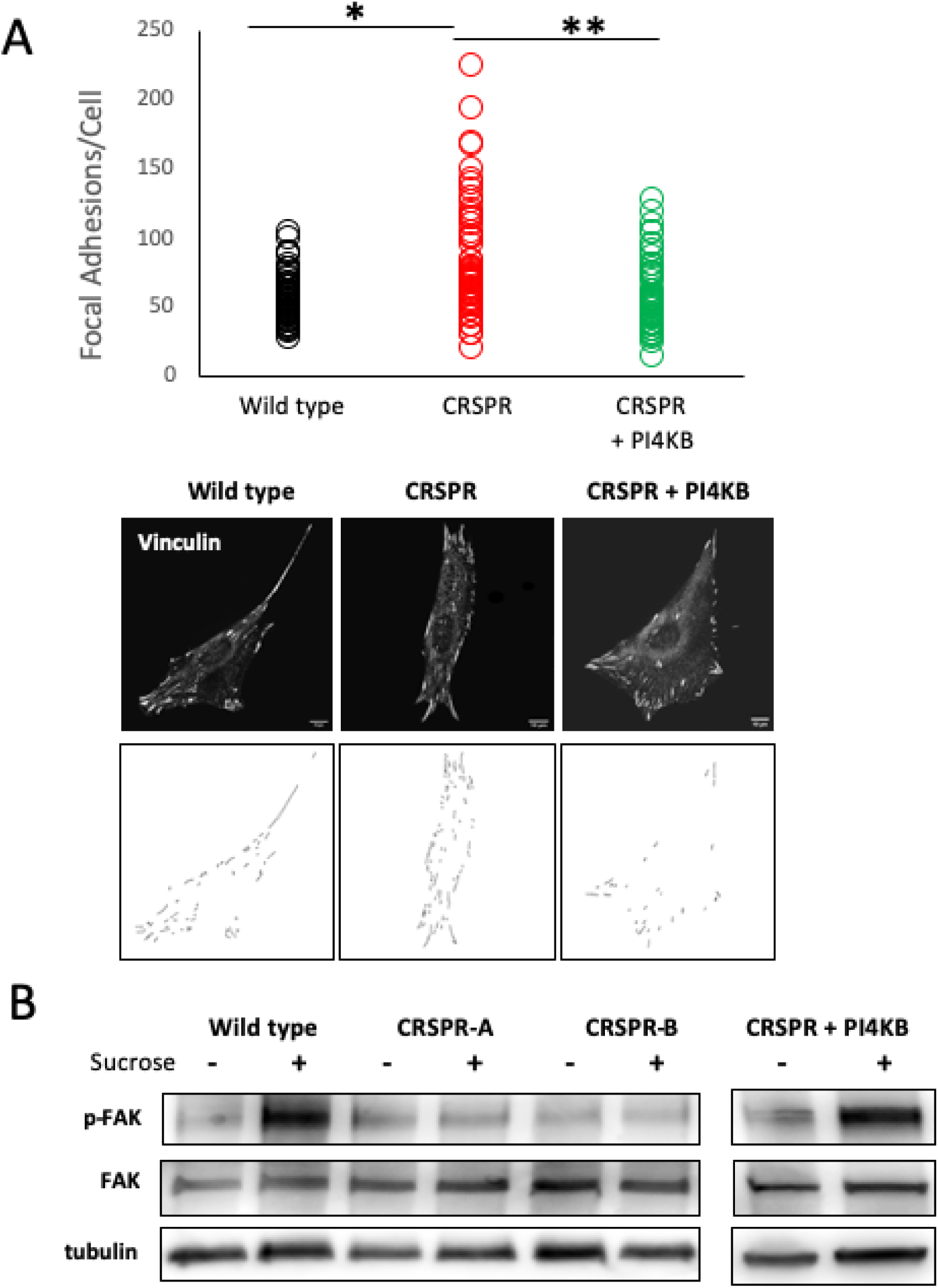
Loss of PI4KIIIβ Increased the number of Focal Adhesions per cell. **A)** The number of FA per cell is shown for wild type cells (n=60), a CRSPR-deleted line (n=59) and for a line rescued with wild type-PI4KIIIβ (n=60). The number of FA in the CRPSR line is significantly greater than in either the wild type line of the one expressing wild type PI4KIIIβ (respectively p< 1 x10^−5^, p<2 × 10^−7^, t-test). Lower images show representative images of FA staining and image segmentation used to count FA for each line. **B)** Blot showing that CRISPR cell lines have a defect in p-FAK phosphorylation in response to 400mM sucrose. This defect is rescued by wt-PI4KB expression.

## DISCUSSION

The central findings of this paper are that the PI4KIIIβ is an important regulator of cell shape, cell migration and FA disassembly. The best known function of PI4KIIIβ is the generation of PI4P from PI (Balla and Balla, 2006; Balla, 2013). In addition, PI4KIIIβ binds to the Rab11a GTPase in a kinase-independent fashion (de Graaf *et al*., 2004; Polevoy *et al*., 2009; Burke *et al*., 2014) and is be involved in Rab11a-dependent control of Akt signaling (Jeganathan *et al*., 2008; Morrow *et al*., 2014) and endosome function (de Graaf *et al*., 2004; Polevoy *et al*., 2009). In our case, a catalytically inactive version of PI4KIIIβ rescues neither the reduced migration nor cell shape defects in PI4KIIIβ-deleted cells. On the other hand, a PI4KIIIβ mutant that has kinase activity but does not bind the Rab11a GTPase (Burke *et al*., 2014) does rescue motility and cell shape distribution to a level equivalent to wild type. This indicates that control of cell migration and shape is independent of Rab11a interaction but requires PI4P generation.

Consistent with an importance for PI4P in migration, our live cell imaging shows directed movement and fusion of PI4P-containing vesicles to the migratory leading edge and to FA. There are also more PI4P vesicles between the nucleus and leading edge than between the nucleus and trailing edge. These vesicles are likely to derive from the Golgi and therefore could be produced by both PI4KIIIβ and PI4KIIβ (Balla, 1998) in the Golgi. PI4P is a precursor for two phosphoinositide’s, PI(4,5)P_2_ and PI(3,4,5)P_3_, that have a well-documented role in cell migration control (Schink *et al*., 2016). Some of the delivered PI4P could be converted to PI(4,5)P_2_ and PI(3,4,5)P_3_ at the leading edge. While PI4P is in large intracellular molar excess over PI(4,5)P_2_ and PI(3,4,5)P_3_, localized pools of PI(4,5)P_2_ in the plasma membrane that regulate ion transport are dependent on an influx of Golgi-derived PI4P (Dickson *et al*., 2014). We propose that pools of PI(4,5)P_2_ and PI(3,4,5)P_3_ regulating cell migration could similarly be dependent on PI4P transport from the Golgi.

Our observation that PI4KIIIβ deletion alone produces defects in cell migration suggests that the PI4P produced by PI4KIIIβ is an important part of cell migration that is not duplicated by PI4KIIβ or other PI4P generating enzymes. Silencing PI4KIIIβ in a breast tumor cell line has previously been shown to attenuate cell migration (Tokuda *et al*., 2014). Here we describe a similar role for PI4KIIIβ in the migration of a non-cancer line. We believe that migratory deficiency in PI4KIIIβ deleted cells is directly related to the changes of cell shape we observe. The majority of wild type NIH3T3 cells display an extended and elongated morphology in culture. On the other hand, a population of PI4KIIIβ deleted cells show fewer elongated cells and more with multiple pseudopodia. We term the multi-pseudopod phenotype “multidirectional” because the cells attempt to move in several directions at once. Elongated cells, on the other hand, generally move along a single axis. The net consequence of the two different types of motion in the different cells is that the multidirectional cells cover smaller distances than elongated ones because they are constantly making small angle turns. Thus, in a wound healing assay, a population of elongated cells will close the wound faster than multidirectional ones. It is important to note that, as a group, both elongated and multidirectional PI4KIIIβ deleted cells have the same migratory parameters as their respective elongated and multidirectional counterparts in wild type cells. They travel at the same velocity and have the same turning behaviors. We propose that deletion of PI4KIIIβ is not impairing the intrinsic migration machinery per se but rather is altering the relative balance of fast-moving and slow moving cells in the overall population.

The increase in FA number we observe in PI4KIII? deleted cells indicates that specific PI4KIIIβ cargo is necessary for normal FA dynamics. FA link the actin cytoskeleton to the ECM. During cell migration, FA are created immediately behind the migratory leading edge and are disassembled at later stages of migration (Parsons *et al*., 2010; Burridge, 2017). Our live-cell imaging indicates that PI4P containing vesicles move to and fuse with FA at both the leading and trailing edges. We favor a model where PI4KIIIβ regulates FA formation by directing the delivery of cargo that initiates FA destruction. One well-characterized process of FA disassembly is the endocytic destruction of a FA via microtubule-mediated delivery of dynamin (Ezratty *et al*., 2005) (Ezratty *et al*., 2009). This activates FA destruction through clathrin-mediated endocytosis. We suggest that delivery of endocytic machinery to sites of FA may be dependent on PI4P-mediated transport. Alternatively, the transport of proteases such as calpain or MMP1 that destroy FA components (Franco *et al*., 2004; Takino *et al*., 2006) could be transported in or on PI4P vesicles.

Another possibility is that PI4P itself, rather than PI4P-dependent cargo, is responsible for FA destruction. Consistent with this idea, depletion of either PI4KIIIβ or IQSec1 results in similar motility impairment and FA disassembly attenuation (D’Souza *et al*., 2020). D’Souza et al propose a model where IQSec1, upon activation by the Ca^2+^ channel Orai1, induces FA disassembly by activating the Arf5 GTPase through the ORP3 lipid exchanger. ORP3 exchanges plasma membrane PI4P for phosphatidyl choline in the ER. Thus, the delivery of PI4P itself could mediate FA destruction through the ORP3/IQSec1 complex.

We favor the idea that the increase in FA number by the loss of PI4KIIIβ is directly responsible for changes in cell shape and migration. The change in FA number is likely responsible for the increase in stress fiber appearance (fig 4C). Stress fibers are long bundles of actin polymers that are typically linked to FA at one or both ends (Burridge and Guilluy, 2016). FA regulate migration because they mediate connection to the ECM and intracellular contractile forces (Parsons *et al*., 2010; Kim and Wirtz, 2013a). While some adhesion is necessary for cell migration, too many FA could impair migration because of excessive adhesion. FA have a reciprocal relationship with cell shape in that the positioning of adhesions determines cell shape but altering the cell shape itself modifies FA location in a cell (Chen *et al*., 2003; Lehnert *et al*., 2004). However, it is a possibility that the changes in cell shape, migration and FA number we observe are regulated by independent PI4KIIIβ and PI4P-dependent pathways.

In summary, we have identified an import role for PI4KIIIβ in cell shape, migration and adhesion. PI4KIII? is likely to be a human oncogene (Waugh, 2012; Morrow *et al*., 2014) and we propose that its role in cancer is related to its regulation of these processes. In the future it will be important to identify the cargo involved.

## ACKNOWLEDGEMENTS

The authors thank Skye McBride and Chloe van Oostende for training and assistance with microscopy and Vera Tang for help with the flow cytometry of the CRSPR lines. We thank the Advanced Imaging Center (AIC) at Janelia Farms for the use of the Bessel Lattice Light Sheet Microscope. We thank Teng-Leong Chew, Satya Khuon, John Hedlleston, Blair Rossetti, and Eric Wait for help with the BLLSM and subsequent image analysis. The PI4KIIIβ-N162A plasmid was a generous gift from Dr. John Burke. FAPP1-GFP was a gift from Tamas Balla. We thank Spencer MacDonald for technical assistance and Redaet Daniel and John Copeland for helpful discussion and critical reading of this manuscript. This work is supported by an operating grant from NSERC (JML).

## METHODS

### Cell lines and culture

The NIH3T3 fibroblast cell line was obtained from ATCC (Manassas, VA, USA). NIH3T3 cells were cultured in Dulbecco’s Modified Eagle Medium (Sigma Aldrich) supplemented with 10% FBS (Thermo Scientific), 1mM sodium pyruvate (Thermo Scientific), 1mM penicillin and streptomycin (Thermo Scientific). Cells are passaged following treatment with 0.25% trypsin protease (GE Healthcare Life Sciences) and counted using TC20 cell counter (Bio-Rad). The PI4KIIIβ targeted RNA sequences for CRISPR deletion of PI4KIIIβ exons 4-5 were 5’-CAGACCGTGTACTCCGAATT-3’, 5’-GGCTCCCTACCTGATCTACG-3’, 5’-ATAAGCTCCCTGCCCGAGTC-3’ (Santa Cruz sc-430739).

### Wound healing and cell tracking

For wound healing, cells were seeded at a density of 1×10^5^ cells/mL in a 24-well ImageLock (EssenBio) plate containing a culture 2-insert well (Ibidi) and incubated a t 37°C at 5% CO_2_. 24hr later, the insert was removed. Images were acquired at a magnification of 10X every 30mins using the IncuCyte ZOOM® Scratch Wound tool (EssenBio). For cell tracking, cells were seeded at 5000 cells/mL in a 24-well ImageLock plate (EssenBio) and images acquired at a magnification of 10mins using the IncuCyte ZOOM (EssenBio). Cells were individually tracked and analyzed in ImageJ.

### Western Blot

Cells were lysed in radioimmunoprecipitation assay buffer (Tris-HCl, pH 7.4, 50mM; NaCl, 150mM; NP-40 1%; sodium deoxycholate, 0.5%; sodium dodecyl sulfate, 0.1%; ethylenediaminetetraacetic acid, 2mM; sodium fluoride, 50mM) supplemented with protease and phosphatase inhibitor cocktails (Roche, Mississauga, Canada). Protein concentrations were determined by Bradford protein assay (Bio-Rad, Mississauga, Canada). Loading buffer was added to 30μg of protein lysate and resolved by SDS-PAGE. The protein was then transferred onto a polyvinylidene difluoride membrane (Millipore, Toronto, Canada) and probed using antibodies for PI4KIIIβ (BD Biosciences 611817; Mississauga, Canada), pan-actin (Cell Signaling Technology 4968), tubulin (1:1000, Cell Signaling Technology 3873; Whitby, Canada), mouse α-FAK (1:200 Thermo Fisher #396500), rabbit α-Phospho-FAK (1:200 ThermoFisher #44-662) as well as anti-rabbit HRP-linked (Cell Signaling Technology catalog no. 7074). Bands were detected with a MicroChemi chemiluminescent system (DNR Bio-Imaging Systems, Toronto, Canada) and intensities were quantified by densitometry using GelQuant (DNR Bio-Imaging Systems).

### Plasmids and Transfections

P4M-GFP and mApple-Talin-N-10 were obtained from Addgene (51469 and 54951 respectively). FAPP-GFP was a gift from Tamas Balla. Cells were grown and transfected on µ-Dish 35 mm, high-wall dishes (Ibidi). For individual P4M and FAPP1 experiments, cells were transfected with 1ug of DNA mixed with Lipofectamine 2000 (Invitrogen) according to manufacturer’s protocol. For P4M and FA imaging, cells were transfected with 1.5μg of P4M, 1.5μg of Talin and 12μL of Lipofectamine 2000 reagent in 250μL.

### Sucrose Stimulation of FAK Phosophorylation

Proliferating NIH3T3 cells and derivatives thereof were serum starved for two hours and then incubated for 10 minutes in serum free DMEM media (Sigma) or stimulated for 10 minutes in serum free media containing 400mM Sucrose (Sigma). Cells were collected and pelleted in ice cold PBS. Cell pellets were lysed in ice cold RIPA buffer containing protease inhibitors (cOmplete, Millipore Sigma #11873580001) and phosphatase inhibitors (PhosSTOP, Millipore Sigma #4906845001) and frozen in 5X LSB.

### Microscopy

For fixed cell microscopy, NIH3T3 cells were seeded on High Precision 1.5H cover glass (Deckglaser) in 12 well cluster plates (Corning). Cells were fixed with 3.7% paraformaldehyde/PBS for 10 minutes, permeabilized for 10 minutes in 0.1% Triton X-100/PBS and incubated for 1 hour in 3% FBS/0.1% Triton X-100/PBS at room temperature. Rabbit α-GM130 (ThermoFisher PA5-95727) and mouse α–tubulin (Cell Signaling 3873S) or γ–tublin (Abcam 27074) were diluted 1:300 in blocking buffer and incubated overnight at 4°C. Mouse α-Vinculin (Millipore Sigma V9131) was diluted 1:300 in blocking buffer and incubated on the cells for two hours (RT). After triplicate PBS washes, Goat anti-Mouse IgG (H+L) Antibody, Alexa Fluor 647 (Thermo Fisher A21235) was diluted 1:500 in PBS and incubated on cells for 1 hour (RT). Cells were mounted on microscope slides (Fisher Scientific 12-544-7) with EverBrite Mounting Medium with DAPI (Biotium 23002) and sealed with CoverGrip Coverslip Sealant (Biotium 23005). Epifluorescent images were acquired with a Zeiss AxioObserver Z1 or a Zeiss AxioObserver M2 microscope with a 63x Plan-Apochromat 1.4 NA oil objective and Zen Blue 2.3 software. Image processing was carried out using ImageJ. The identification of FA immunostained for vinculin were processed as described by Utku Horzum et. al.(Horzum *et al*., 2014)

### Live Cell Microscopy

Time-lapse series of cells in phenol-free DMEM were recorded at 37°C on a Zeiss LSM880-AxioObserver Z1 microscope equipped with 63X Plan Aprochromat (NA 1.4) oil objective. We used an AiryScan detector in FAST mode and the bandpass emission filter 495-620nm. Live-cell time lapse series were also captured using a Bessel Beam Lattice Light Sheet Microscope equipped with detection objective (Semrock; NA of 0.54). A 5mm coverslip was loaded to a piezo stage and submerged in a 37°C water bath. Time lapse series of 175 Z-planes and 350 volumes were captured at 63X magnification with S Piezo Offset 50, and an interval of 0.4 µm. Raw data were assembled into .tif hyperstacks using ImageJ. 3D renders were then generated for analysis using Imaris.

## Supplementary File Information

**Supplementary Video 1**. Migrating NIH3T3 fibroblast transfected with FAPP1 to visualize PI4P vesicles.

**Supplementary Video 2**. Migrating NIH3T3 fibroblast transfected with P4M to visualize PI4P vesicles.

**Supplementary Video 3**. Migrating NIH3T3 fibroblast transfected with P4M to visualize delivery and fusion of PI4P vesicles with the plasma membrane during migration.

**Supplementary Video 4**. Migrating NIH3T3 fibroblast transfected with FAPP1 to visualize delivery and fusion of PI4P vesicles with the plasma membrane during migration.

**Supplementary Video 5**. Migrating NIH3T3 fibroblast transfected with P4M to visualize the distribution of PI4P vesicles in the cytoplasm.

**Supplementary Video 6**. Non-migrating NIH3T3 fibroblast transfected with P4M to visualize the distribution of PI4P vesicles in the cytoplasm.

**Supplementary Video 7**. Migrating NIH3T3 fibroblast transfected with FAPP1 to visualize the distribution of PI4P vesicles in the cytoplasm.

**Supplementary Video 8**. Non-migrating NIH3T3 fibroblast transfected with FAPP1 to visualize the distribution of PI4P vesicles in the cytoplasm.

**Supplementary Video 9**. Migrating NIH3T3 fibroblast transfected with GFP-P4M and mApple-Talin to simultaneously visualize FAand PI4P vesicles. Video shows the migratory leading edge.

**Supplementary Video 10**. Migrating NIH3T3 fibroblast transfected with GFP-P4M and mApple-Talin to simultaneously visualize FA and PI4P vesicles. Video shows the migratory trailing edge.

## Notes

**Funding:** This work was supported by operating funds from the Natural Sciences and Engineering Research Council of Canada (JML).

### Competing Interest Statement

The authors have declared no competing interest.

